# S1 represents multisensory contexts and somatotopic locations within and outside the bounds of the cortical homunculus

**DOI:** 10.1101/2022.08.29.505313

**Authors:** Isabelle A. Rosenthal, Luke Bashford, Spencer Kellis, Kelsie Pejsa, Brian Lee, Charles Liu, Richard A. Andersen

**Author notes:** Correspondence (I.A.R.).

## Abstract

The responsiveness of primary somatosensory cortex (S1) to physical tactile stimuli is well documented but the extent to which it is modulated by vision is unresolved. Additionally, recent literature has suggested that tactile events are represented in S1 in a more complex, generalized manner than its long-established topographic organization. To better characterize S1 function, neural activity was recorded from a tetraplegic patient implanted with microelectrode arrays in S1 during 1s stroking touches to the forearm (evoking numb sensation) or finger (naturalistic sensation). Touch conditions included visually observed first person physical touches, physical touches without vision, and visual touches without physical contact which occurred either to a third person, an inanimate object, or the patient’s own body in virtual reality. Two major findings emerged from this dataset. The first was that vision strongly modulates S1 activity, but only if there is a physical element to the touch, suggesting that passive observation of touches is not sufficient to recruit S1 neurons. The second was that despite the location of the recording arrays in a putative arm area of S1, neural activity was able to represent both arm and finger touches in physical touch conditions. Arm touches were encoded more strongly and specifically, supporting the idea that S1 encodes tactile events primarily through its topographic organization, as well as in a more general manner encompassing larger areas of the body.

## Introduction

The sense of touch is important for implementing dexterous, adaptable action plans [1]–[5] and creating a sense of ownership and agency over one’s body [6]–[8]. The primary source of information for tactile sensations is input from peripheral mechanoreceptors, but multisensory integration [9], [10] plays a role as well, especially visual information [11]–[14].

Primary somatosensory cortex (S1) is one of the first cortical areas to receive incoming tactile information, relayed from the spinal cord through the thalamus [15]. S1’s responsiveness to physical touch and its topographic organization have been extensively documented [16]–[20], but the extent to which multisensory information is represented in S1 has yet to be resolved. If S1 functions mostly as a simple relay between the thalamus and higher order cortical areas such as S2 and posterior parietal cortex, it might not be activated by visual information, even if it is relevant to tactile experiences; alternatively, S1 could be a site of early cortical processing for multisensory integration. A large body of literature addressing this precise question has found that S1 responds to observed touch when it occurs in others but not oneself [21]–[28]. However, a significant number of studies have failed to find evidence of this phenomenon [29]–[32].

A similar but distinct question concerns whether S1 is modulated by vision when it is paired with a physical touch event. Psychophysically, the visual enhancement of touch has been well-established: tactile acuity is enhanced when a touched area is observed, even when the visual input is non-informative [14], [33]–[36]. EEG experiments have shown that combining visual and tactile stimuli modulates the P50 somatosensory evoked potential, which is thought to originate in S1 [37]–[41]. MEG studies have suggested that the topographic mapping of fingers shifts in S1 based on the relative timing of visual and tactile signals [42], [43]. Transcranial magnetic stimulation (TMS) over S1 negatively affects the ability to detect or discriminate touches, if the accompanying visual information incorporates a human hand rather than a neutral object [44]–[46]. Thus biologically relevant visual information appears to be used as a predictive signal and modulate S1 encodings of tactile events.

Like its role in integrating multisensory stimuli, S1’s topographic organization appears to be more nuanced than first thought. Recent experiments suggest that although S1 maintains a gross topographic representation of the body as laid out in the earliest human cortical stimulation studies and confirmed many times since [16]–[20], [47], it also contains other more complex levels of tactile representation [48]–[50]. Studies of the primate hand have shown that S1 neural activity contains non-linear interactions when multiple fingers are stimulated simultaneously [48], [50], supporting the idea that S1 carries information beyond a linear report of inputs from tactile receptors. In humans, S1 has recently been shown to represent body parts outside of their traditionally defined areas [51], [52].

To interrogate S1’s representations of touch across body locations and multisensory contexts, electrophysiological recordings in a human tetraplegic patient with two microelectrode arrays (Blackrock Microsystems, Salt Lake City, UT) implanted in the putative area 1 of the S1 arm region [53] were collected. The patient retained enough tactile ability after spinal cord injury to sense short stroking stimuli delivered to his arm and finger. Touch conditions occurred on either the patient’s arm, finger, or an inanimate object, in a variety of multisensory contexts (**Table 1**). Our results provide evidence that tactile information in S1 is encoded as part of the well-established cortical homunculus as well as in a more general manner which encompasses larger areas of the body. Additionally, we find that S1 does not respond to observed touches to oneself, another person, or an object, but that vision does modulate neural activity when it is paired with physical tactile stimulation. This finding suggests that passively observing visual information depicting touches fails to meet some threshold of relevance or attention necessary to activate S1 neurons.

**Table 1:** Touch modality conditions tested, each presented with the color code representing that touch type throughout the paper. Each modality with a physical touch stimulus is coded with ‘*’. All modalities with a stimulus on the arm end in ‘a;’ all modalities with a stimulus on the finger end in ‘f.’ Touches were single strokes delivered by an experimenter using a pressure-sensing rod; in *BL and VrFP trials the participant wore a Vive Pro Eye headset.

| Modality | Location | Touch Type | Description |
| --- | --- | --- | --- |
| *FPa | Arm (a) | *FP | The participant observed, in first person (FP) perspective, a real physical touch to his body. |
| *FPf | Finger (f) |  |  |
| *BLa | Arm | *BL | The participant fixated on a non-informative dot in virtual reality, effectively blindfolding (BL) him during a real physical touch to his body. |
| *BLf | Finger |  |  |
| VrFPa | Arm | VrFP | The participant observed, via virtual reality and in first person perspective (VrFP), a touch to his body without any physical contact. |
| VrFPf | Finger |  |  |
| TPa | Arm | TP | The participant observed, in third person (TP) perspective, a real physical touch to another person's body. |
| TPf | Finger |  |  |
| Obj | Object | Obj | The participant observed a real physical touch to an inanimate object (Obj), a wooden block. |

## Methods

### Participant

A C5-level incomplete tetraplegic participant (male, 32 years old) was recruited and consented for a brain-machine interface (BMI) clinical trial including intracortical recording and stimulation. The participant was implanted with microelectrode arrays in three locations in the left hemisphere: the supra-marginal gyrus (SMG), ventral premotor cortex (PMv), and primary somatosensory cortex (S1). This paper only examines data in S1, which was recorded using two 48-channel 1.5mm SIROF-tipped (sputtered iridium oxide film) microelectrode arrays (Blackrock Microsystems, Salt Lake City, UT). Given the curvature of sensorimotor cortex and the need to implant arrays on the gyral surface, it is likely the S1 micro-electrode arrays are located in Brodmann area 1 (BA 1). Additional details pertaining to the arrays and the specifics of surgical planning are described in [53]. At the beginning of data collection, the participant was 6.5 years post-injury and 5 years post-implant. All procedures were approved by the Institutional Review Boards (IRB) of the California Institute of Technology, University of Southern California, and Rancho Los Amigos National Rehabilitation Hospital.

### Experimental Paradigm

Two anatomical locations were examined across a set of tactile and visual conditions. The two locations selected were a “finger” location on the back of the thumb where the participant reported naturalistic sensations, and an “arm” location near the back of the elbow where the participant reported numb sensation. These locations were selected on the basis of a preliminary mapping of the participant’s tactile capabilities on the arm and hand using Semmes-Weinstein filaments at varying strengths, which took place two days prior to the first experimental session.

Although the different task conditions (**Table 1**) did not all include both a physical and a visual component, all employed the same style of touch: a 1-second stroke over approximately 6cm of skin. The touch was delivered by a plastic rod (or a virtual facsimile of one), built in-house, which had a raised button on one end (1.5 × 2 cm) that was passed along the touch location. The rod housed a load cell which was used to record the pressure applied and align the onset of touch to neural recordings.

Each trial consisted of an inter-trial-interval (ITI) of 5s with an additional 0-3s jitter, followed by a 1s touch stimulus and 1s post-touch phase. In trials with a visual component, the visual component began approximately 0.5s before touch onset (for instance, in several conditions the experimenter could be seen to move the pressure-sensing rod towards the touch location prior to contact with the touch location, **Fig 1a**). A total of 11 conditions were examined, incorporating 6 touch types and 3 touch locations (**Table 1**).

**Figure 1:**
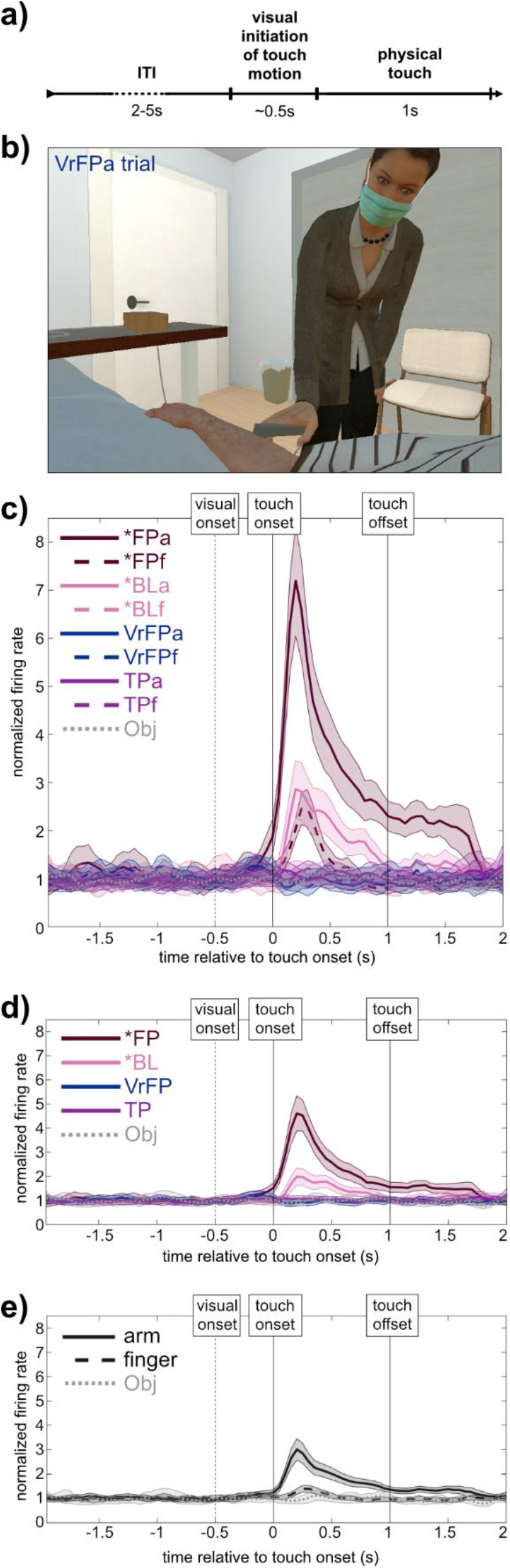
**a)** Task time course. Visual initiation of touch motion was only perceived by the participant in touch types with visual content (*FP, VrFP, TP, Obj). **b)** Sample frame from a VrFPa trial, presented using a virtual reality headset. **c)** Example smoothed firing rate of one S1 channel to each tested modality of touch (n=70 trials/modality). Shaded area surrounding each line indicates standard error of the mean (SEM). **d)** and **e)** depict activity of the same channel averaged across modalities to isolate touch type and effector respectively (i.e. *FPa and *FPf in **(c)** are averaged to yield *FP in **(d)**.

### Data Collection

Neural data was recorded from each microelectrode array using a 128-channel Neural Signal Processor (Blackrock Microsystems) as 30,000 Hz broadband signals. Data was collected in 8 sessions over 6 months, in two sets (see **Table 1** for task condition descriptions). In the first set, the participant observed real physical touches to his body in first person (*FP), the same touches delivered to someone else (third person; TP), and touches to an inanimate object (Obj). This set was collected over the first two months in 4 sessions. In the second set, the participant experienced real physical touches without visual touch information (blind; *BL), and saw touches being delivered to him in first person using virtual reality, without any physical touch component (VrFP). The second set was collected over the third to sixth months in 4 sessions.

Within a session, data was collected in series of 11-trial runs. Each run contained 10 trials of the same condition and one catch trial. Within the two sets, runs were pseudorandomly shuffled so there were no two runs of the same condition back to back in any session. 70 trials (7 runs) were collected in every condition.

The conditions in set two required a virtual reality headset; a Vive Pro Eye was used to display a virtual environment run with Unity, which closely mimicked the data collection room and gave the participant a first-person perspective over a virtual body with a size, gender, and posture reflecting his own body. In the virtual environment, a virtual experimenter was animated to deliver touches in a manner resembling the real experimenter (**Fig. 1b**). The human avatar for the virtual experimenter was taken from the Microsoft Rocketbox Avatar Library (https://github.com/microsoft/Microsoft-Rocketbox/). For *BL conditions, the headset was used as a blindfold, and displayed a non-informative white dot in the center of a black field of view which the participant was instructed to fixate on.

Although outside the scope of this paper, additional conditions were collected along with the ones analyzed here. In set one, conditions in which the participant imagined the touches being delivered without any external tactile stimuli were also obtained. In set two, a touch type identical to VrFP except that the participant’s virtual body was composed of abstract blocks rather than a realistic human body was collected, and a condition in which the participant viewed an inanimate object being touched in virtual reality was also acquired.

### Preprocessing and Temporal Alignment of Neural Data

Firing rates for each electrode were extracted in 50ms bins from the broadband signal in a multi-unit, unsorted fashion [54], [55], using a threshold of −3.5 times the noise RMS of the continuous signal voltage. This multi-unit channel activity was aligned within each trial to the physical or virtual moment of contact between the touch sensor and the item being touched (i.e. touch onset). In conditions with a physical touch component, touch onset was calculated using the pressure readings obtained from the rod used to deliver touches; in conditions with only a virtual touch component, touch onset was calculated using the timing of Unity animations.

To normalize firing rates, within each run and each channel, a mean baseline firing rate was calculated from the time period 4s to 2.5s prior to each touch onset and averaged across trials. The firing rates of each channel at every time point were divided by this baseline.

### Decoding Analysis

Linear Discriminant Analysis (LDA) pairwise classifiers were used to probe the linearly decodable information within and across task conditions (**Figs. 2a, 3**). Normalized firing rate data was binned into either 0.5s or 0.1s bins, depending on the analysis. Within each bin, data was randomly split equally into train/test partitions. This split occurred 1000 times and was balanced each time to include equal numbers of trials from every condition tested (70 trials per condition = 35 trials each in train and test).

**Figure 2:**
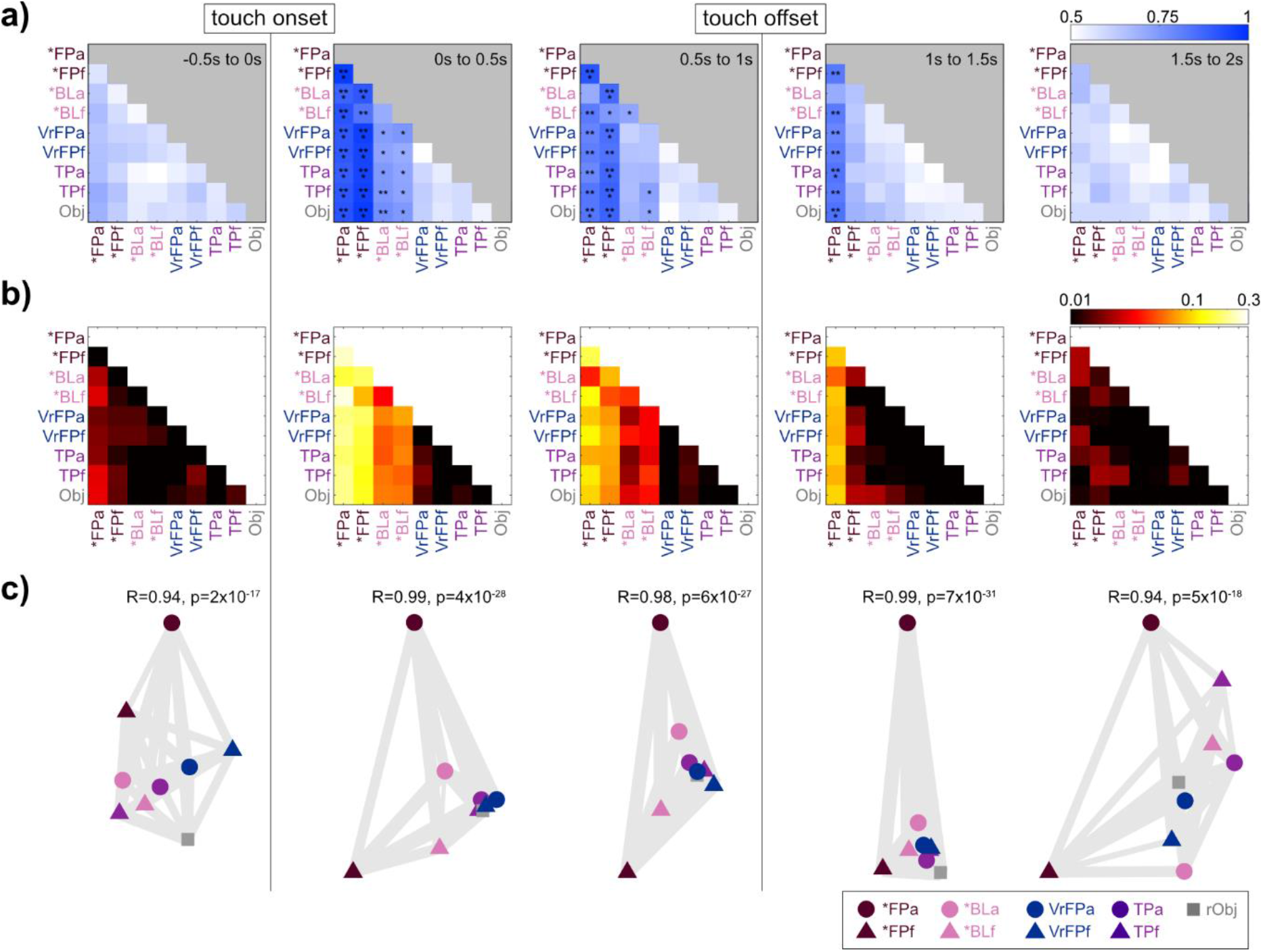
**a)** Pairwise identity decoding results. At each 0.5s time bin, a LDA classifier was trained to distinguish between each pair of modalities based on the top 40 principle components of multiunit activity. The 70 trials per modality were randomly divided in half to generate train and test data 1000 times, and the accuracies of the resulting decoders were averaged together to yield the values in the confusion matrices. Asterisks represent significantly different accuracies relative to a null distribution which was generated by training the same decoder on data with shuffled labels 1000 times. * = significantly different 95% confidence intervals (CIs); ** = 97.5% CIs; *** = 99% CIs. **b)** Representational similarity analysis was performed on touch-onset-aligned multi-unit S1 channel activity, and resulting representational dissimilarity matrices (RDMs) are shown. Distances between conditions (plotted on log axis) are cross-validated Mahalanobis distance with multivariate noise correction; a distance of 0 indicated conditions are statistically indistinguishable. **c)** Multi-dimensional scaling (MDS) plots of RDMs in **(b)**. Axes are arbitrary but have been rotated for consistency across time bins. Gray lines between condition icons are “rubber bands” whose thickness is based on the goodness of fit of the scaling. A relatively thinner, more “stretched” band between conditions indicates that in a plot that fully captures neural geometry, the conditions would be closer together.

**Figure 3:**
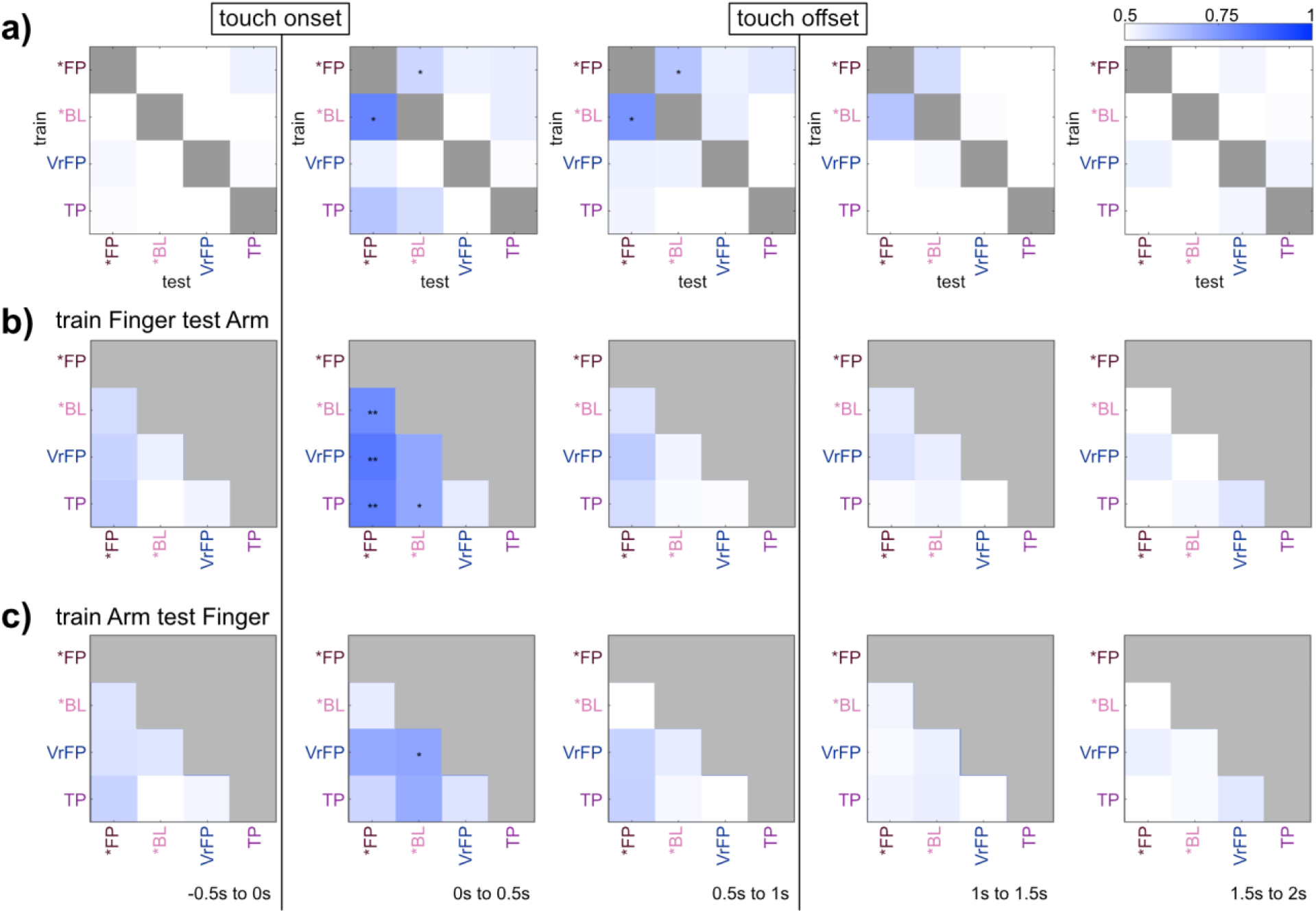
Generalization decoding results. **a)** Effector was decoded across all pairs of touch types. For example the bottom right square of each grid represents the average accuracy when a decoder is trained to distinguish Arm v Finger on TP trials and tested on *FP trials. **b)** Pairs of touch types were decoded, training on Finger trials and testing on Arm trials. For example the bottom right square of each grid represents the average accuracy when a decoder is trained to distinguish TP vs *FP on Finger trials and tested on Arm trials. **c)** The same procedure as in **(b)** except that training occurred on Arm trials and testing on Finger trials. All decoders used 140 trials in training and testing respectively, 70 of each effector. All statistics and plotting conventions as in **Fig. 2a**.

Singular value decomposition (SVD) was used to perform dimensionality reduction on the initial 96 multi-unit channels of the training dataset. Average firing rate data from both train and test datasets in each bin was projected on the top 40 features capturing the most variance in the training data.

LDA classifiers were fit to the resulting data using MATLAB’s *fitcdiscr* function across the 1000 iterations. The overall performance of each classifier was taken as the average performance and 95% confidence intervals on this estimate were taken from the distribution of accuracies across iterations. This analysis was repeated on a null dataset in which condition labels were shuffled across trials in order to generate chance-level performance of the classifier. Significance was calculated by comparing the accuracy percentile values of the classifiers with their null counterparts.

### RSA and MDS

Representational Similarity Analysis (RSA) was employed on normalized firing rate data to assess the relationships between touch conditions (**Fig. 2b,c)**[56], [57]. Cross-validated Mahalanobis distance with multivariate noise normalization was used as the measure of dissimilarity [58]. The noise covariance matrix was estimated from the data and regularized towards a diagonal matrix to ensure that it would be invertible. The cross-validated Mahalanobis distance is an unbiased measure of square Mahalanobis distance with the added benefit of having a meaningful zero-point [58], [59]. The larger the Mahalanobis distance between two conditions, the more discriminable their neural patterns. If the patterns are fully indiscriminable, their distance is 0. This continuous measure is directly related to discrete classification performance with pairwise LDA. Cross-validated Mahalanobis distance is thus less affected by common activation patterns across conditions in comparison to other measures such as Pearson correlation. The python package *rsatoolbox* (https://github.com/rsagroup/rsatoolbox) was used to compute noise covariance and generate representational dissimilarity matrices (RDMs).

Data was cross-validated across 7 splits, divided by the 10-trial runs the data was originally collected in, and RDMs were generated independently on data divided into 0.5s bins. The resulting RDMs were symmetric across the diagonal, with meaningless values on the diagonal itself.

RDMs were visualized with multi-dimensional scaling (MDS) using the MATLAB toolbox *rsatoolbox* (https://github.com/rsagroup/rsatoolbox_matlab)[57]. MDS allows for distances in RDMs to be visualized intuitively in a lower-dimensional space while preserving these distances as much as possible. The MDS visualizations used a metric stress criterion to arrange conditions without assuming any category structure a priori. The stress is visualized on MDS plots (**Fig. 2c**) in the form of grey “rubber bands” stretched between points – the thinner the band, the more the true distances between points are distorted by the low dimensional MDS mapping to be further apart than in the high dimensional RDM.

### Tuning and Onset Analysis

Tuning properties of multi-unit channels were assessed via linear regression analysis. In each 500ms bin corresponding to 1s before touch onset to 2s after touch onset, normalized firing rates for each channel were fit to a linear regression model based on the following equation:

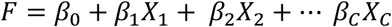

where F = vector of firing rates on each trial, X = one-hot-encoded matrix signaling condition identity for each trial, *β*= estimated regression coefficients indicating level of tuning to each condition, and C = number of conditions tested. In addition to data from every trial, F also included 70 entries (to match the number of trials per condition), corresponding to *β*_0_, containing the baseline firing rate of the channel across all trials. This baseline was calculated as a mean of channel activity 4s to 2.5s before touch onset in every trial.

For each channel and condition fit with linear regression, a student’s t-test was performed to assess the null hypothesis *β*= 0. If the null hypothesis was rejected, the channel was determined to be tuned to that condition in comparison to its baseline firing rate. P values were corrected for multiple comparisons using the Bonferroni-Holm method within each channel.

A bootstrap analysis was run for 1000 iterations, in which all conditions were randomly sampled with replacement to yield 70 trials each, to assess significant differences in numbers of tuned channels across conditions (**Fig. 4**).

**Figure 4:**
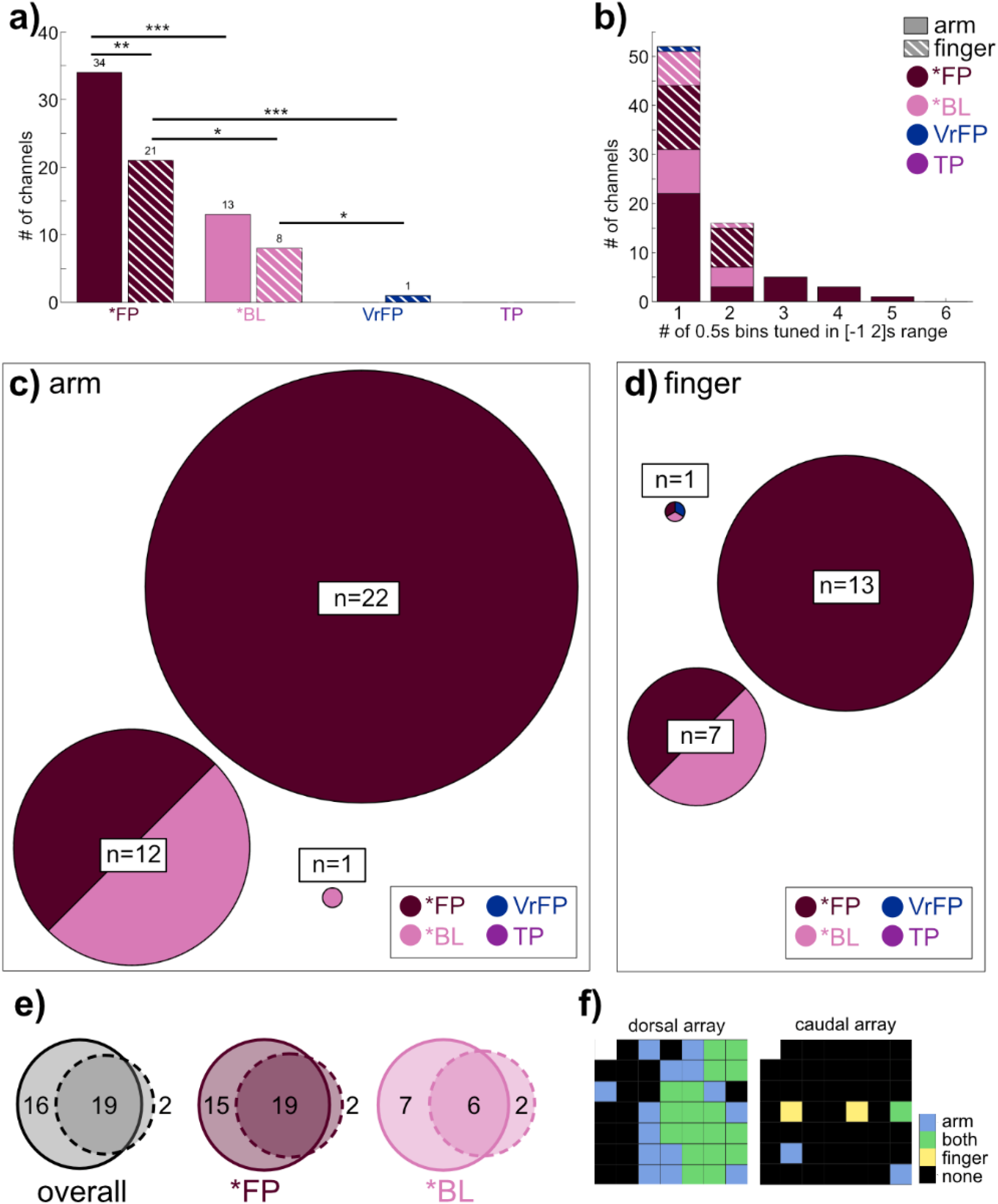
Channels selective for any touch modality (p<0.05, Bonferroni-corrected linear regression analysis) at any time bin in the −1 to 2s range relative to touch onset were examined. **a)** Total number of channels tuned within arm and finger touch conditions. Asterisks indicate non-overlapping 95% CIs. **b)** Histogram indicating the range of time that channels were tuned. Tuning was performed in 0.5s non-overlapping time bins (maximum bins a channel could be tuned to in the −1 to 2s range was 6. **c)** Circles indicate the specific set of modalities each channel was tuned to, within arm touch modalities, and have a diameter proportional to the number n channels tuned to that set. **d)** Plotting as in **(c)** but based on finger touch conditions. **e)** Distribution of channels tuned to arm (solid line circle), finger (dashed line circle), or both, across all touch conditions (left) or within touch types (middle, right). **f)** Array map of implanted electrodes, indicating locational tuning across all conditions.

The channels identified as tuned to any condition in the period of 1s before touch onset to 2s after touch onset were analyzed to determine the average timing onsets and offsets of their tuned responses. Within a condition, firing rates of all tuned channels were averaged together in 50ms bins, and the 95^th^ percentile of the distribution of average baseline firing rates was computed. The onset time for the condition was the middle of the first time bin in which the firing rate rose above the 95^th^ percentile of the average baseline. The offset time was calculated as the middle of the first time bin in which the firing rate dipped below the 95^th^ percentile of the average baseline, after onset. 95% confidence intervals were constructed for the onset and offset times by bootstrapping over trials within the tuned channels 10,000 times.

## Results

S1 responses to visual and tactile stroking stimuli along the arm and finger in a human tetraplegic participant were recorded via two intracortical microelectrode arrays. Within arm and finger locations, neural responses to four touch types were examined **(Table 1; Fig 1**). A fifth touch type (Obj) used an inanimate object as a control rather than a body location, resulting in a total of 9 conditions across locations and touch types. 70 trials were collected in each condition.

Multi-unit channel activity recorded during these trials was aligned to the physical or virtual moment of contact between the touch sensor and the item being touched (touch onset). In visual conditions (*FP, VrFP, TP, Obj), visual information predicting the touch was available beginning approximately 0.5s before touch onset, as the experimenter could be seen beginning the motion towards the touch target (**Fig. 1a**). In totality, the task comprised 9 conditions (**Table 1**); the average firing rate of a single channel to each condition is plotted as an example (**Fig. 1c**). The task was designed such that data could be averaged across location (**Fig. 1d**) or averaged across touch type (**Fig. 1e**) to better isolate neural responses to these factors.

### Condition identity decoding

Linear discriminant analysis (LDA) was performed on the top 40 dimensions of the multi-unit channel data, sub-selected over 1000 train/test divisions for equal class sizes and averaged together (**Fig 2a**). Classifiers were trained on every pair of conditions, using average firing rates binned in 0.5s increments. No significant decoding occurred prior to touch onset. In the first 0.5s following touch onset, conditions containing a physical touch (*FP, *BL) could be meaningfully distinguished from purely visual conditions (VrFP, TP, Obj) in all cases, and could be significantly distinguished from one another in every case except *BLa vs *BLf (accuracy = 70% [60-80%]), which only became significant 0.5s later (72% [61-81%]). *FPa vs *BLa was highly decodable with an accuracy of 87.7% [80-94.3%] despite the two classes only varying on the basis of visual information; similarly, *FPf vs *BLf obtained an accuracy of 83.7% [74.3-91.4%].

Overall in the first time bin post touch onset, *FPa and *FPf were highly distinguishable from other conditions, especially those without physical touch. *BLa and *BLf were less significantly distinguishable. In the following time bin (0.5-1s), this relative disparity in classification accuracies remained true, but classifications were overall weaker (pairwise one-sided t-test across all accuracies, p=3×10^−9^).

In the [1-1.5s] bin which occurred immediately post touch offset, *FPa was the only condition that could be distinguished from the other conditions. 1.5s post touch onset, no classifiers obtained significant decoding accuracy.

To examine decoding on a finer time scale, the same LDA classifiers as described above were run in touch-onset aligned 0.1s bins (**Supplemental video 1**). No significant decoding occurred before the 0-0.1s bin. In this bin, *FPa could be significant decoded from all conditions apart from *FPf and *BLa, but no other classifiers were significant. In the following 100ms (0.1-0.2s post touch onset) *FP/*BL could be significantly distinguished from all other conditions with the exception of *FPf vs *BLf. By 0.3-0.4s post touch onset, decoding was overall weaker than in this first bin suggesting the time period of 0-0.2s following touch onset contains the strongest touch representations. By 0.7-0.8s post touch onset, all classifiers ceased to be significantly accurate.

### Representational Similarity Analysis (RSA)

To better visualize the relationships between different task conditions, RSA [56] was used on the same multi-unit activity as analyzed with the linear classifier (**Fig 2b**). Representational dissimilarity matrices (RDMs) were computed based on the cross-validated Mahalanobis distance with multivariate noise correction [58]. For visualization purposes, multi-dimensional scaling (MDS) was used to scale the relationships captured in the RDMs into two dimensions (**Fig 2c**) [57].

There was a high level of similarity between pairwise decoding (**Fig 2a**) and the RDMs (**Fig 2b**) during touch encoding (for the three consecutive time bins post touch onset, r>0.89; p<1×10^−12^ in all cases, Bonferroni corrected). This similarity is expected since both methods assess the discriminability of neural activity averaged across conditions.

In the 0.5s prior to touch onset, distances between conditions formed a pattern which shared a mild correlation with activity post touch onset; activity was most correlated between the −0.5-0s RDM and the 1-1.5s RDM (**Fig 2b**, Pearson correlations between −0.5-0s RDM and RDMs 0-2s, in chronological order: r = 0.67, 0.61, 0.70, 0.38; p = 3×10^−11^, 3×10^−9^, 1×10^−12^, 5×10^−4^, Bonferroni corrected).

Once touch occurred, the initial RDM within 0-0.5s post touch onset contained a strong pattern that remained stable during the touch and afterwards, although it became weaker as time elapsed (Pearson correlations between 0-0.5s RDM and RDMs 0.5-2s, in chronological order: r = 0.96, 0.86, 0.62; p = 2×10^−43^, 1×10^−24^, 9×10^−10^, Bonferroni corrected).

Within the 0-0.5s bin, touch types with only visual stimuli (VrFP, TP, Obj) were less distinguishable and tightly grouped together, while the physical touch types (*FP, *BL) were more distinct from one another and therefore more spread out (two-sample t-test on distances within VrFP/TP/Obj vs distances within *FP/*vBL: p=9×10^−5^).

*FP and *BL varied in their level of separation from the non-physical touch types (**Fig 2c**, 0-0.5s bin). *FP (mean=0.17; std=0.03) had more distance to the non-physical touch types than *BL (mean=0.04; std=0.03). These sets of distances were significantly different from each other (paired t-test, p=3×10^−5^). Additionally, during and immediately after the touch, *FP/*BL arm representations were grouped distinctly from the finger representations: on the MDS plots (**Fig 2c**), arm conditions are consistently grouped separately (above) finger conditions.

### Location and touch type generalization decoding

To investigate if body location information generalizes across touch types, LDA classifiers were trained to differentiate arm/finger conditions within one touch type and tested on another (**Fig 3a**). During the touch (0-1s), body location information generalized within physical touch conditions; it was possible to train the classifier on *FP and decode body location from *BL, or vice versa. The strongest decoding was achieved in the 0-0.5s bin by the decoder that trained on *BL and tested on *FP (accuracy = 79.6% [68.6-90%]). Post touch offset, generalization was no longer possible.

When the same decoding problem was performed in 0.1s bins (**Supplemental video 2**), the only significant accuracies were at 0.1-0.2s post touch onset for the same subset of conditions significant in the 0-0.5s bin, indicating a small window of time when body location can generalize strongly across touch types. The opposite question was also interrogated: can a classifier trained on one body location successfully decode the type of touch presented using another body location? In this case, for both classifiers that trained on finger data and tested on arm data (**Fig 3b**), and classifiers that trained on arm data and tested on finger data (**Fig 3c**), the only significantly decodable instances occurred in the 0-0.5s time bin.

Decoding was notably asymmetric between training on finger/testing on arm and the opposite paradigm: *FP could be strongly distinguished from all other conditions when training on finger and testing on arm, but could be significantly distinguished from no conditions when training on arm and testing on finger. Additionally, there were smaller asymmetries in significance across the two paradigms when distinguishing *BL from VrFP or TP. Decoding in 0.1s time bins was overall weaker, and only was significant in the 0.2-0.3s time bin for training on finger, testing on arm (**Supplemental video 3**), while training on arm, testing on finger never reached significance at any time point (**Supplemental video 4**).

### Individual channel tuning analysis

To investigate the tuning properties of individual channels within the S1 arrays, linear regression analysis was performed in 0.5s bins aligned to touch onset. The most channels were tuned to *FPa (34 [95% CI=30, 45] out of 96 channels total), a number significantly greater than the number of channels tuned to *FPf (21 [18, 26]) or *BLa (13 [10, 22]; **Fig. 4a**). *FPf elicited more tuned channels than *BLf, which trailed at 8 [6, 16] channels. Of the non-physical touch conditions, only 1 [0, 4] channel was tuned to VrFPf. The number of time bins that tuned channels were responsive to a given condition was quantified (**Fig. 4b**). No tuning to finger conditions occurred for longer than 2 bins (1s). *BLa tuning followed the same rule, but *FPa trials elicited up to 5 bins (2.5s) of responsive activity.

The overlap across tuned arm conditions within channels was calculated (**Fig 4d**). 22 channels were tuned to *FPa solely, while 12 channels were tuned to *FPa and *BLa together. Similarly, 13 channels were tuned to *FPf solely, while 7 channels were tuned to *FPf and *BLf together. In finger conditions (**Fig. 4d**), fewer channels were tuned overall. Within arm and finger, channels were nearly all tuned to *FP conditions (**Fig. 4c, d**).

The overlap of tuned channels across all arm and finger conditions was determined (**Fig 4e**). Overall, most channels were tuned to both arm and finger (n=19), while 16 channels were only tuned to arm conditions. Only 2 channels were tuned to solely finger conditions. Within touch types, a similar pattern emerged. In *FP, 19 channels were tuned to both arm and finger, 15 to just arm, and 2 channels to just finger. In *BL, 6 channels were tuned to arm and finger, 7 to just arm, and 2 to just finger. Lastly, the position of tuned channels across all arm and finger conditions was plotted on diagrams of the microelectrode arrays (**Fig 4f**). Channels tuned to both arm and finger were clustered together, surrounded by channels tuned to arm only. The two exclusively finger-tuned channels were located on the opposite array from the other channels.

The average tuned response curves of channels tuned to *FP and *BL conditions were examined (**Fig. 5**), calculated as deviation from the distribution of baseline activity: *FPa onset = −25ms [95% CI= −75, 75]; *FPf onset = 125ms [75, 125]; *BLa onset = 25ms [25, 75]; *BLf onset = 75ms [75, 125]. The average offset times of tuned activity were also determined, relative to onset of the touch stimulus: *FPa offset = 1125ms [575, 1575]; *FPf offset = 825ms [625, 875]; *BLa offset = 875ms [575, 925]; *BLf offset = 875ms [525, 925].

**Figure 5:**
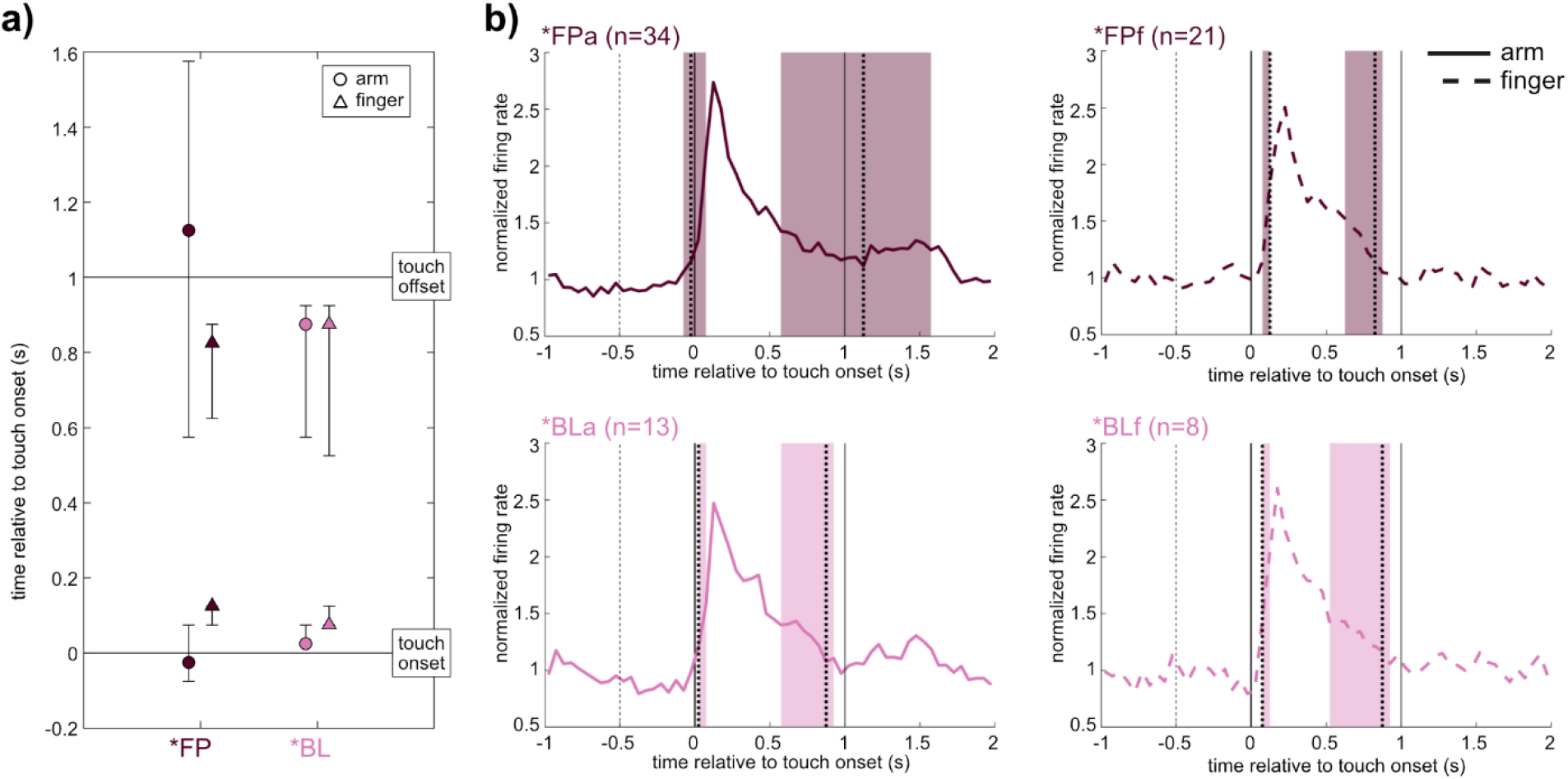
**a)** Onset and offset of channel tuning. Onset of tuning was calculated based on the average normalized firing rate across all tuned channels and condition trials, as the first time bin where average activity exceeded the 95^th^ percentile of the distribution of average baseline responses. Offset was calculated on the same data as the first time bin to dip below the 95 ^th^ percentile of baseline activity after the peak firing rate. Error bars represent 95% CI obtained by bootstrapping 10,000 times over trials. **b)** Average responses to individual conditions in tuned channels, relative to touch onset (vertical black line at 0s); touch offset is plotted as a vertical black line at 1s. Vertical dotted lines indicates onset and offset of response with colored background depicting 95% CIs. n=number of tuned channels.

While there was substantial variance across trials and electrodes, a major trend emerged from this analysis. Within *FP and *BL, arm onset times always occurred before finger onset times, while offset times were similar across conditions with the exception of the wide variance of *FPa (**Fig. 5a**). Within arm and within finger, onsets times were not statistically different although the mean of *FPa onsets occurred slightly prior to touch onset. In all conditions (**Fig. 5b**), activity peaked sharply immediately following touch onset, followed by a gradual decrement of activity back to baseline.

## Discussion

To examine how S1 represents tactile events based on their location and their multisensory context, electrophysiology data from the putative area 1 of the S1 arm region were examined. Within arm and finger locations, touch types varying in their tactile and visual content were tested (**Table 1**). It is worth noting that the participant’s long-term tactile impairment could have resulted in representations of touch in S1 that are altered relative to healthy humans. This is unlikely to be a major effect on the findings of this study, because recent work has shown that topographic representations in S1 are highly preserved in tetraplegic people, even years post-injury [60], [61].

Analysis of this rich, exploratory dataset suggested two main conclusions about local neural activity where the multi-electrode arrays were implanted: 1) This S1 area is specialized for arm representations but is capable of representing touch information from the finger in a more general manner; 2) This S1 area is modulated by vision during physical touches, but is not activated by vision on its own.

### Neural activity is specialized for arm touches and represents finger touches more generally

Immediately after touch onset in FP* and BL*, arm conditions were separated from finger conditions, based on neural activity visualized using MDS (**Fig. 2c**). This division based on touch location continued until the touch ceased. *FP and *BL were both separable by touch location, although *BL was significantly less separable. This pattern was also evident in the linear decoding analysis, where *FPa vs *FPf was immediately highly decodable upon touch onset, while *BLa vs *BLf was only significantly decodable one time step (0.5s) later (**Fig. 2a**). *FPa could be distinguished from all other conditions for much longer than any other condition – up to 1.5s post touch onset – and was the first condition to become decodable in the 0.1s bin classifier (**Supplementary video 1**).

Despite *FP seeming to contain more robust location information than *BL, classifiers trained on either *FP or *BL and tested on the other to distinguish arm from finger trials achieved significant decoding (**Fig. 3a**). Locational information was therefore sufficiently present in *BL conditions to allow for generalization to and from *FP conditions. Through analysis of the 0.1s bin decoders, it appears the bulk of this locational information was present 0.1-0.2s post touch onset.

The tuning analysis demonstrated that while arm and finger touches were both represented in S1, the representations were not equal. More channels overall were tuned to arm than finger conditions (**Fig 4a**), and *FPa trials elicited tuning for up to 5 time bins (2.5s) while *FPf tuning only lasted up to two time bins (1s; **Fig. 4b**). The vast majority of tuned channels were either tuned to solely arm conditions or both arm and finger conditions (**Fig. 4e**). Only two channels were tuned to solely finger conditions, suggesting the bulk of the neural population recorded is not selective to finger touches specifically, but may be activated by body touches more generally in addition to arm touches specifically. *FP and *BL each contained roughly equivalent numbers of channels tuned to solely arm or to both finger and arm, and very few channels tuned solely to finger (**Fig. 4e**). Onset analysis of *FP and *BL revealed a trend that appeared to mesh with this pattern – *FPa onset occurred 0.15s before *FPf onset, and *BLa onset occurred 0.05s before *BLf (**Fig. 5a**). In other words, arm conditions elicited sharply tuned neural responses that began before the tuned responses to finger conditions (**Fig. 5b**).

To summarize, neural activity elicited by physical touches delivered to the arm formed patterns distinct from the activity elicited by touches to the finger. Individual channels tended to be tuned to both arm and finger, or just arm conditions, but rarely just finger conditions. Tuned activity started earlier for arm conditions than finger conditions. This evidence builds a picture of a region of S1 which is primarily geared towards representing arm touches. A neural sub-population of this region is also is capable of representing finger touches, albeit less strongly or specifically.

There could be several reasons for this difference between the two tested locations. One is that, due to the spinal cord injury, the participant was able to sense one location more strongly and naturalistically than the other. The patient reported finger sensations to be more natural, yet neural finger representations were weaker. This makes the participant’s uneven tactile impairment an unlikely culprit for the differences in location encoding.

If the differences between arm and finger representations are not primarily due to differences in spinal nerve damage and tactile impairment, then they are likely due to differences in the neural representations of these locations. The distribution of tuned channels appeared geographically distributed when mapped to the implanted micro-electrode arrays – the upper array contained the bulk of activity, with a nucleus of channels tuned to both arm and finger conditions and a surrounding of channels tuned solely to arm conditions (**Fig. 4f**). The only two finger-specific channels were located on the lower array. Cortical curvature may have resulted in electrodes recording from varying cortical layers within S1.

From prior work with this participant, it is known that intracortical microstimulation (ICMS) of the S1 arrays studied here elicits cutaneous and proprioceptive sensations primarily in the arm, with a much smaller number of sensations in the fingers [53]. It is likely the arrays, especially the dorsal array (**Fig. 4f**), are located in S1’s arm region. The neural response to finger touches detailed here contribute to the growing literature suggesting that although S1 does maintain a gross representation of the body along the lines of the homunculus laid out in the earliest human cortical stimulation studies and confirmed many times since [16]–[20], it also contains other more complex levels of tactile representations [48]– [51]. Most recently, Muret et al. (2022) used MRI to show that different body locations are represented in S1 in areas beyond their primary topographic area. Our findings support and expand this finding, indicating that S1 encodes highly specific and rapid responses to touches through its established topography, but tactile information from much larger anatomical areas may activate S1 in a more general manner.

### Visual information modulates neural activity if accompanied by a physical stimulus

S1 neural activity was restricted to conditions which contained a physical tactile stimulus, and less than 2% of channels were tuned to visual-only conditions **(Fig. 4a**). Although several variations on visual touches without physical stimuli were tested (VrFP, TP, Obj), they were not represented in a discriminable manner from one another in S1 (**Fig 2**). VrFP and TP did not elicit representations of touch location information in S1 activity, whether decoded in an identity or generalization problem (**Fig 2a, 3a**). Across all methods in this study, there was no detectable encoding of tactile information in S1 from purely visual stimuli.

In contrast, *FP trials contained visual information paired with a physical stimulus and, immediately after touch onset, they could be easily discriminated from all non-physical conditions and from *BL trials which contained the same physical stimulus minus the visual content (**Fig 2a**). The strong performance of *FPa vs *BLa and *FPf vs *BLf classifiers indicate the presence of visual information is sufficient to change the touch encoding in S1. Visual information also appeared to affect the length of time a touch representation occurred in S1, as *FPa was decodable for much longer than any other condition.

RSA demonstrated that the pattern of responses immediately prior to touch onset was mildly correlated with activity during the touch itself, suggesting there is some effect of a visual approach of a tactile stimulus before an expected touch occurs (**Fig 2b**). However, a much stronger stable pattern of activity emerged once the touch actually began, as indicated by the correlations between the RDM of the first 0.5s to the following RDMs. This relationship is evident in the MDS plots generated based on neural activity (**Fig 2c**). In the second following touch onset, *FP and *BL conditions are separated from all other touch types and from each other. In particular, *FPa and *FPf are highly dissociated from the other conditions. The presence of visual information generalized across touch location to some extent – a classifier trained on finger trials could distinguish *FP vs *BL in arm trials, but not vice versa (**Fig. 3 b,c**). The ability to decode visual information in a manner than generalizes across body location also appeared to be present quite late relative to touch onset; the 0.1s bin decoder only achieved any significance in the 0.2-0.3s time bin relative to touch onset. These findings speak to a more general distinction between visual and blind physical touches existing in S1 finger touches, which is overridden by more specific information in arm touches that are not able to generalize to other body parts.

There were large populations of channels tuned to *FPa and *FPf, and within these populations there were sub-groups also tuned to *BLa and *BLf respectively (**Fig 4c, d**). Blind and visual touches appear to activate the same population of neurons, but touches with a visual component activate additional neurons on top of this population.

These results suggest that visual information is enough to distinguish two otherwise identical physical touches in S1, but visual information on its own, whether it relates to oneself (VrFP), another person (TP) or an inanimate object (Obj) is not sufficient to engage S1. This finding is especially intriguing because while it is clear that visual information affects experiences of touch [11], [36], [62], a rapidly evolving scientific literature is still deciding the role of vision in modulating S1 [21], [25], [26], [28]–[31].

To our knowledge, the results presented here are the first to examine the effect of vision on S1 using human electrophysiology, which as employed here is highly localized to a sub-region of S1, with high spatial resolution of spiking activity. The bulk of prior literature has used fMRI, MEG, and EEG to address this question, data which capture whole brain dynamics at a relatively low spatial resolution, and likely include membrane potentials that do not produce spikes. Across these experiments, results have varied, finding either that S1 either has no response to observed touch [29]–[31] or does respond to observed touch [21], [24]–[26], [28]. One notable difference between the studies that uncovered seemingly contradictory findings is the type of task employed.

The majority of experiments finding S1 modulation to observed touches employed a touch-relevant task during data collection, whether it be counting touches [23], [28], answering qualitative questions about the touch type [22], [24], [25], or rating touch intensity [21]. The experiments which found no effect of observed touch on S1 tended to employ either non-touch-related tasks [29] or simply asked participants to passively observe the stimuli [30], [31]. Thus it is possible that a relevance threshold, modulated by higher order brain areas, must be exceeded in order for S1 to represent observed touches [40]. If this is true, the fact that S1 did not respond to visual stimuli when the participant passively observed touches in this study agrees with the existing literature, despite the differences in data types. This effect could also explain why visual information does modulate tactile representations of physical touch – the physical component of the touch activates S1 as it would in a blind touch, but additionally higher order areas integrate the visual input as sufficiently relevant to the tactile input such that vision affects S1 simultaneously.

What might be the role of this modulation? It is known that vision modulates experiences of touch in a variety of ways, including effects like visual enhancement of touch [14], [33]–[36] in which non-informative vision of a body part improves tactile perception. Our results suggest that touch-relevant visual information elicits an earlier tuned response over more neurons, and results in a representation of touches that are highly distinguishable in terms of location and multisensory content. All of these attributes have the potential to contribute to visual enhancement of touch. Indeed, these results agree with prior literature which has suggested that S1 is modulated by paired visual and tactile stimuli [37]– [41], [43], [63], [64], and has also shown that S1 is a necessary component of body-centered visuotactile integration [44]–[46] and reflects predictions of tactile events from visual signals [65].

S1 can be modulated by concepts as high-level as affective significance, as was shown in a study which examined the effect of perceived gender of a person delivering a caress to heterosexual men [62]. S1 is also affected by motor planning, presumably expecting the sensory consequences of upcoming actions [66], and by imagining touch sensations [67], [68]. It is likely that when S1 is modulated by visual information, it is not directly interacting with the visual system but instead affected by upstream areas which are implementing some version of a forward model to determine expected tactile inputs.

### Conclusion

This study represents a broad exploration of how different types and locations of touch affect a small area in the putative arm region of S1. It contributes to the growing body of literature suggesting that S1 contains a large-scale, highly specific topographic organization, with more generalized touch representation throughout. We also find that visual information depicting touches, either to oneself, to another person, or to an object, are not sufficient to activate S1 in a measurable way. However, a blindly sensed physical touch and a visually seen physical touch are represented distinguishably in S1– both elicit strong responses that share commonalities, such as how touch location is encoded, but they are not identical.

Taken as a whole, these findings demonstrate that S1 contains a nuanced and complex encoding of tactile experiences that is to some degree multisensory. Future endeavors should aim to examine these same conditions in a larger population of individuals, both healthy and with a variety of levels of sensorimotor impairment. There are many practical applications for a better understanding of S1, including the improvement of restorative devices seeking to artificially generate tactile sensations in deafferented limbs and prosthetics [53], [69]–[71]. By better understanding how naturalistic tactile sensations are encoded in S1, and how they interact with cues from other sensory modalities, we can improve our ability to generate biomimetic artificial stimuli.

## Acknowledgements

We thank S. Wandelt and D. Bjånes for helpful discussions and insights, S. Norman for invaluable assistance building the pressure-sensing equipment, and participant FG for his effort and dedication to the study.

## Author Contributions

I.A.R., L.B., S.K., and R.A.A. designed the study. I.A.R. developed the experimental tasks. I.A.R. and L.B. collected data. I.A.R. analyzed the results. I.A.R. and L.B. interpreted the results. I.A.R. wrote the paper. I.A.R., L.B., and R.A.A. reviewed and edited the paper. K.P. coordinated regulatory requirements of clinical trials. C.L. and B.L. performed the surgery to implant the microelectrode arrays.

